# Comprehensive cell surface proteomics defines markers of classical, intermediate and non-classical monocytes

**DOI:** 10.1101/2020.02.20.958009

**Authors:** Benjamin J. Ravenhill, Lior Soday, Jack Houghton, Robin Antrobus, Michael P. Weekes

**Affiliations:** Cambridge Institute for Medical Research, University of Cambridge, Hills Road, Cambridge, CB2 0XY, UK

## Abstract

Monocytes are a critical component of the cellular innate immune system, and can be subdivided into classical, intermediate and non-classical subsets on the basis of surface CD14 and CD16 expression. Classical monocytes play the canonical role of phagocytosis, and account for the majority of circulating cells. Intermediate and non-classical cells are known to exhibit varying levels of phagocytosis and cytokine secretion, and are differentially expanded in certain pathological states. Characterisation of cell surface proteins expressed by each subset is informative not only to improve understanding of phenotype, but may also provide biological insights into function. Here we use highly multiplexed Tandem-Mass-Tag (TMT)-based mass spectrometry with selective cell surface biotinylation to characterise the classical monocyte surface proteome, then interrogate the phenotypic differences between each monocyte subset to identify novel protein markers.

## Introduction

Monocytes play critical roles in the response to infection and inflammation and antigen presentation. Key effector functions are mediated by differentiated cells, including macrophages and myeloid dendritic cells. Monocytes can be divided into subsets based on the presence of the cell surface lipopolysaccharide (LPS) co-receptor CD14, and Fc-Gamma Receptor III (CD16). CD14++ CD16-classical monocytes account for ∼85% of circulating monocytes; intermediate cells (CD14++ CD16+) for ∼5% and non-classical cells (CD14+ CD16++) for ∼10% of circulating monocytes^1, 2^. It has become increasingly well established that each subset plays divergent roles in different diseases, as well as differing in the ability to secrete cytokines and respond to pathogen associated molecular patterns (PAMPs). Classical monocytes are phagocytic and readily secrete inflammatory cytokines. Conversely, the CD16 positive non-classical cells are poorly phagocytic and are suggested to secrete TNFα in response to some stimuli, but less of other pro-inflammatory molecules^2,3, 4^. Intermediate monocytes are increased in diseases such as severe asthma, rheumatoid arthritis and sarcoidosis, and there is some evidence for expansion of classical monocytes in atherosclerosis^5-8^. It is still unclear whether intermediate monocytes represent a truly distinct monocyte subset, or merely a transitional stage between classical and non-classical cells^9^.

Cells of the innate and adaptive immune systems can be categorised on the basis of microscopic appearance and expression of plasma membrane (PM) proteins, enabling separation by fluorescence activated cell sorting (FACS). As such, systematic evaluation of the entire cell surface proteome expressed by a given immune cell population is a powerful tool to characterise cellular function and distinguish cell types. Previous studies of monocytes subsets have exclusively examined transcriptional differences and offer varying depths of information about subset markers, suitability of individual proteins for discriminating subsets by cell surface flow cytometry and the importance of each protein in different pathological states^2, 8, 10, 11^. Usage of multiple complementary grouping systems has the benefit of improving cellular assignment, in addition to enabling discovery of new cellular phenotypes^8,12^.

Here we use selective surface protein biotinylation with multiplexed tandem mass tag (TMT)-based mass spectrometry to directly measure the first comprehensive surface proteome of primary human classical monocytes from three donors. 373 classical monocyte cell surface proteins were quantified, and the relative abundance of each protein estimated. The surface proteome of classical, intermediate and non-classical cells was then compared to identify unique markers of each subset. Amongst these, Integrin alpha subunit 5 (ITGA5), complement receptor 1 (CR1/CD35) and Leukotriene B4 receptor (LTB4R) were defined as markers of classical monocytes, and Sialic Acid Binding Ig Like Lectin 10 (SIGLEC10) as a marker of non-classical cells.

## RESULTS

### Establishing a definitive classical monocyte surface proteome

To establish a surface proteome map, monocytes were enriched from three independent peripheral blood mononuclear cell (PBMC) donations from healthy donors by negative selection with magnetic beads. Classical monocytes were then enriched by FACS after staining for CD86, CD14 and CD16 **(Supplementary Fig. S1)**. Selective surface oxidation and aminoxybiotinylation was used to label cell surface glycoproteins, which were enriched from cellular lysates using streptavidin beads, then digested into peptides using trypsin. Peptides were labelled with TMT, samples combined and then quantified by MS3 mass spectrometry^13, 14^ (**Fig. 1a**).

**Figure 1.**
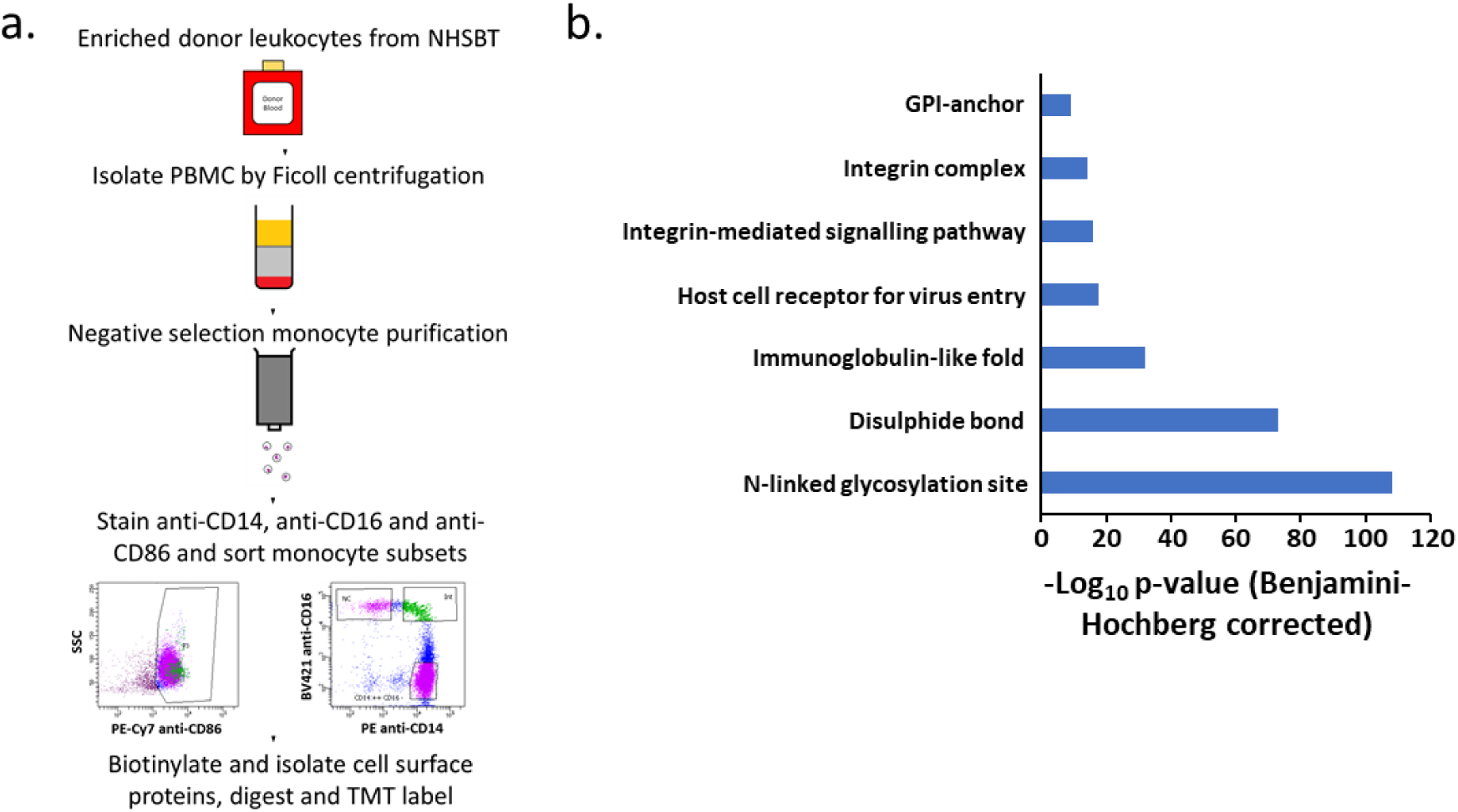
Overview of experimental strategy and protein enrichment scores. a) A schematic overview of the experimental approach. b) Enrichment of pathways within all 373 classical monocyte surface proteins in comparison to all human proteins as background, using DAVID software. Benjamini-Hochberg adjusted p-values are shown for each pathway.

437 proteins were identified from the three samples, of which 373 were annotated ‘cell surface’, ‘plasma membrane’, ‘extracellular’ or with a short Gene Ontology term as previously described^13, 14^ (**Supplementary table S1d**). Application of DAVID software to determine which pathways were enriched amongst these proteins indicated the presence of multiple components of integrin complexes and cell-cell junctions in addition to ‘glycosylation’ and ‘disulphide bond’, serving to validate our selective labelling approach (**Fig. 1b**).

### Quantification of the classical monocyte cell surface proteome

We used a method derived from identity based absolute quantitation (iBAQ) to compare the contribution of each protein to the classical monocyte cell surface proteome. The summed MS1 maximum precursor intensity for each protein across all matching peptides was divided by the theoretical number of tryptic peptides 7-30 amino acids in length. Values thus effectively represent an average across three donors, offering the opportunity to provide precise information on the overall abundance of each PM protein, independent of individual genetic variation. Abundance values ranging over approximately five orders of magnitude were found, with 21 proteins collectively contributing 68.6% of the cell surface proteome, whilst individually contributing >1% (**Fig. 2**). The five most abundant surface proteins, CD44, SPN, ICAM3, ITGB2 and BSG accounted for ∼25% of the surface proteome, with CD14 representing ∼3.5% (**Fig. 2**). The summed abundance of class I Major Histocompatability antigen (MHC) accounted for 2.0% of surface proteins, and class II MHC 0.7%.

**Figure 2.**
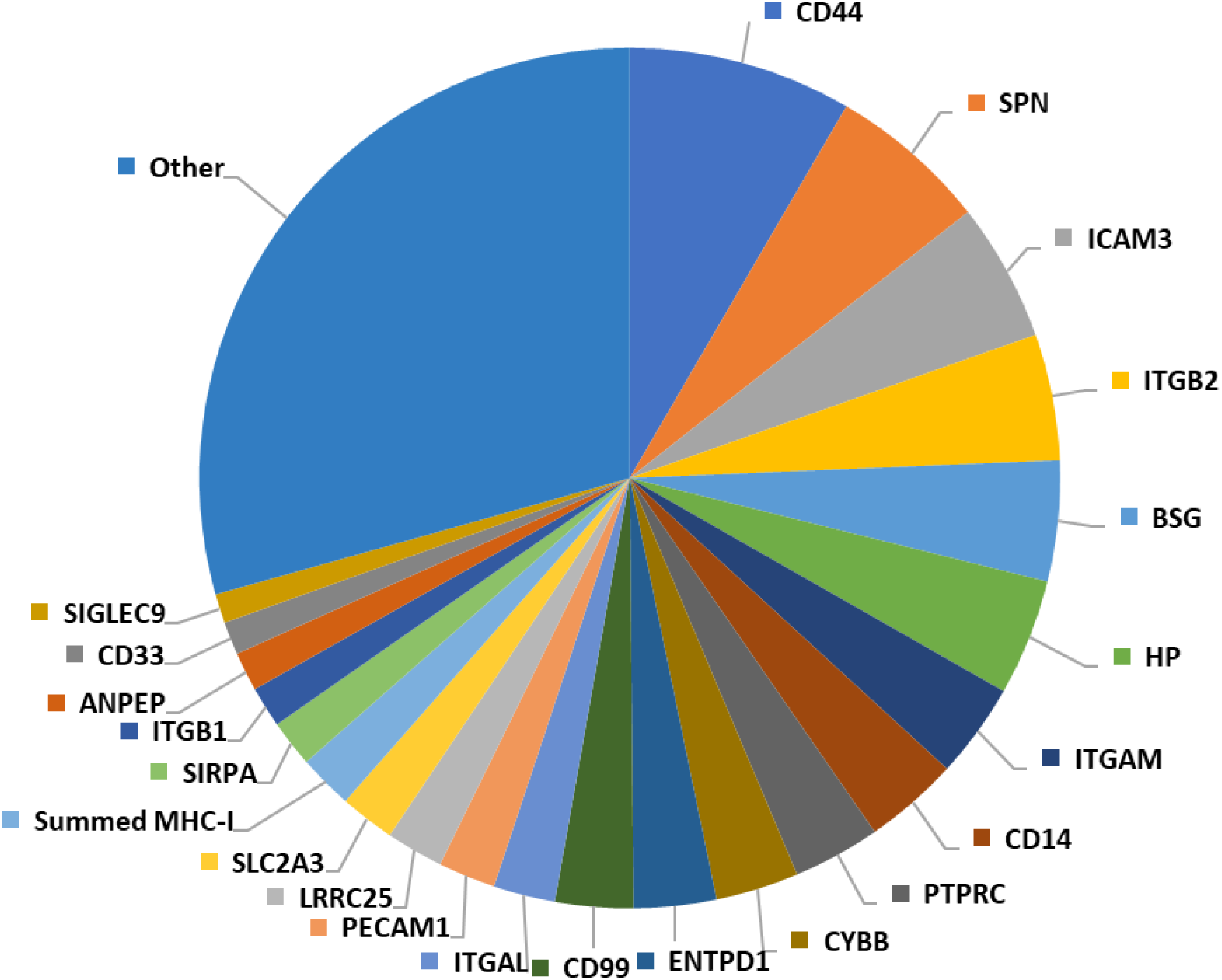
Pie chart showing the relative contribution of individual proteins to the classical monocyte surface proteome. Proteins contributing <1% are included in the ‘other’ category. The summed abundances of all MHC class I molecules and MHC class II molecules were considered in order to account for different alleles expressed by individual donors.

By using multiplexed TMT-based proteomics, this study offered the opportunity to directly measure the variability in expression of whole surface proteomes across different donors. 76.1% of proteins exhibited a <30% coefficient of variation (%CV). There was also no systematic inverse relationship between abundance and variability, i.e. less abundant proteins were not systematically more poorly quantified (**Supplementary Fig. S2**).

### Comparison of the cell surface proteome of different monocyte subsets

Although the majority of circulating monocytes belong to the classical subset, the non-classical and intermediate subsets play an increasingly appreciated role in different diseases. Selective cell surface biotinylation with MS3-based mass spectrometry was used to investigate phenotypic differences between each subset in biological triplicate. From this analysis, 313 proteins annotated ‘cell surface’, ‘plasma membrane’, ‘extracellular’ or with a short GO term were quantified (**Supplementary table S1e**). Principle component analysis (PCA) suggested that classical monocytes had a more distinct cell surface proteome in comparison to non-classical and intermediate cells (**Fig. 3a**). K-means clustering with 1-20 classes suggested there were at least eight distinct patterns of protein expression between monocyte subsets (**Fig. 3b-c**). Examples of proteins selectively enriched on classical monocytes (Cluster A) included CD99 antigen and Sialic Acid-Binding Ig-Like Lectin 3 (CD33), validating previous transcriptomic studies^2, 10^. Clustering also highlighted a number of proteins enriched in non-classical cells (Cluster C), including CD16a (FCGRIIIA), the previously reported non-classical monocyte marker SIGLEC10^2^ and Tetraspanin 14 (TSPAN14), which had not previously been reported. Distinguishing intermediate monocytes from other subsets was more challenging, however several candidate cell surface markers were identified including Solute Carrier Family 6 Member 6 (SLC6A6) (Cluster B). We then calculated the Benjamini-Hochberg corrected p-values for each monocyte subset comparison. This confirmed that the ‘classical’ and ‘intermediate’ clusters were enriched in proteins showing significantly differential abundance between the subsets (**Supplementary table S1e**). The enrichment for the third ‘non-classical’ cluster was poor, in keeping with the appearance of the k-means analysis.

**Figure 3.**
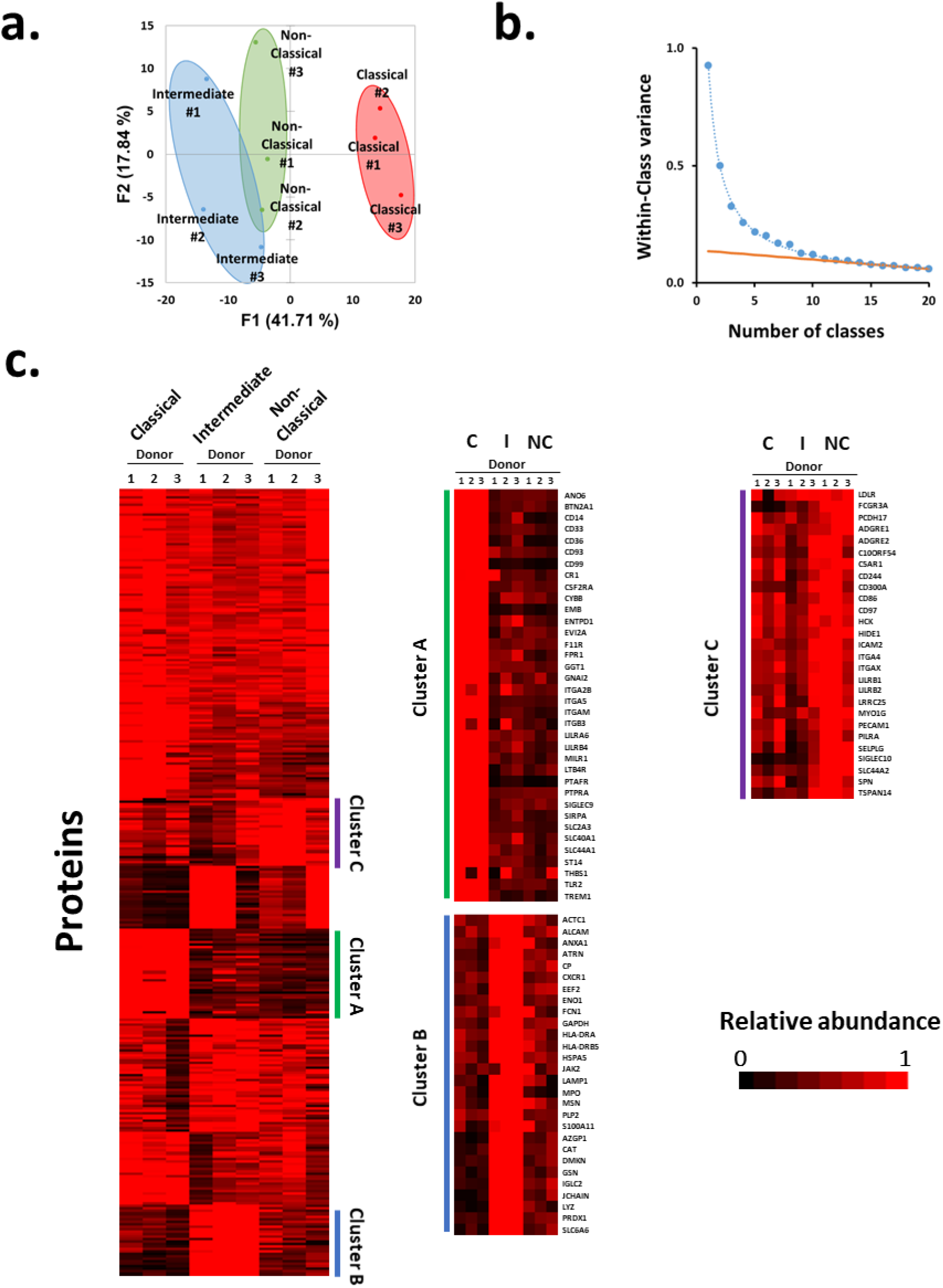
Analysis of the surface proteome of monocyte subsets. (a) Principal component analysis of all 313 proteins annotated ‘cell surface’, ‘plasma membrane’, ‘extracellular’ or with a short GO term. The grouping of the samples suggested that subset type was the major driver of variability as opposed to donor-specific differences. (b) The number of distinct classes of protein expression between monocyte subsets plotted against within-class variance. Proteins were clustered using a k-means approach with 1-20 classes, and the summed distance of each protein from its cluster centroid was determined. This summed distance becomes smaller as more clusters are added, but the rate of decline decreases with each iteration, eventually reaching a fairly steady rate of decline (orange line) that is reflective of overfitting. Clusters added before this point reflect the underlying structure in the protein data, whereas clusters subsequently added through overfitting add no additional useful information. For these data the point of inflexion was between eight and nine classes, suggesting that there are at least eight distinct surface protein profiles. (c) K-means based hierarchical cluster analysis of the 313 proteins identified at the surface of the different classes of monocyte. Right panels – enlargements of clusters particularly enriched in proteins of one class (the adjacent bar colour indicates where each enlargement matches the original cluster plot). C – Classical monocyte sample, I – Intermediate monocyte sample, NC – Non-classical monocyte sample.

### Comparison to RNA microarray data

Previous analyses of monocyte subsets have mostly been performed at the level of RNA expression^2, 8, 10, 11^. Hierarchical clustering was used to assess the complementarity between one of the most comprehensive previous transcriptomic studies^2^ and this proteomic analysis. For each protein, each data type was normalised to a maximum of 1 after averaging signal:noise (proteomics) or median fluorescent intensity values (microarray) across donors (**Supplementary Fig. S3**). RNA and protein data for the classical subset clustered separately from non-classical and intermediate subsets. However, intermediate protein-RNA did not neatly separate from the non-classical pair suggesting that these cells may be more phenotypically similar to one another than classical monocytes, in keeping with the PCA (**Fig. 3a**). Another recently published transcriptomic analysis^15^ used single-cell RNAseq to analyse Lin-HLA-DR+ index sorted cells. These included dendritic cells, monocytes and a population of contaminating NK cells. A phenograph-clustering algorithm aimed to identify within each cluster (i) cell types and (ii) differentially-expressed genes (DEGs). Eight clusters were identified, and clusters 1 and 3 were related most closely to classical monocytes (CD14^hi^ CD16^-^, Figure 3B from Duterte et al^15^), with cluster 7 most closely related to mixed intermediate and non-classical monocytes (CD16^+^, Cluster 7 in the same figure). Comparison of DEGs to our cell surface protein-level data largely confirmed that these markers were differentially expressed at both the level of protein and RNA within classical and non-classical monocytes (**Supplementary Fig. S4**).

### Validation of cell surface subset markers

Flow cytometry was used to validate a group of monocyte subset markers identified by proteomics, using independent samples from three different healthy donors of European heritage. SIGLEC10 validated as a marker most abundantly expressed by non-classical monocytes (**Fig. 4a-b**)^2^. ITGA5, CR1 and LTB4R also validated as markers of classical monocytes identified by proteomic data (**Fig. 4a-b**), in line with previous high throughput analyses for ITGA5 and LTB4R ^2^.

**Figure 4.**
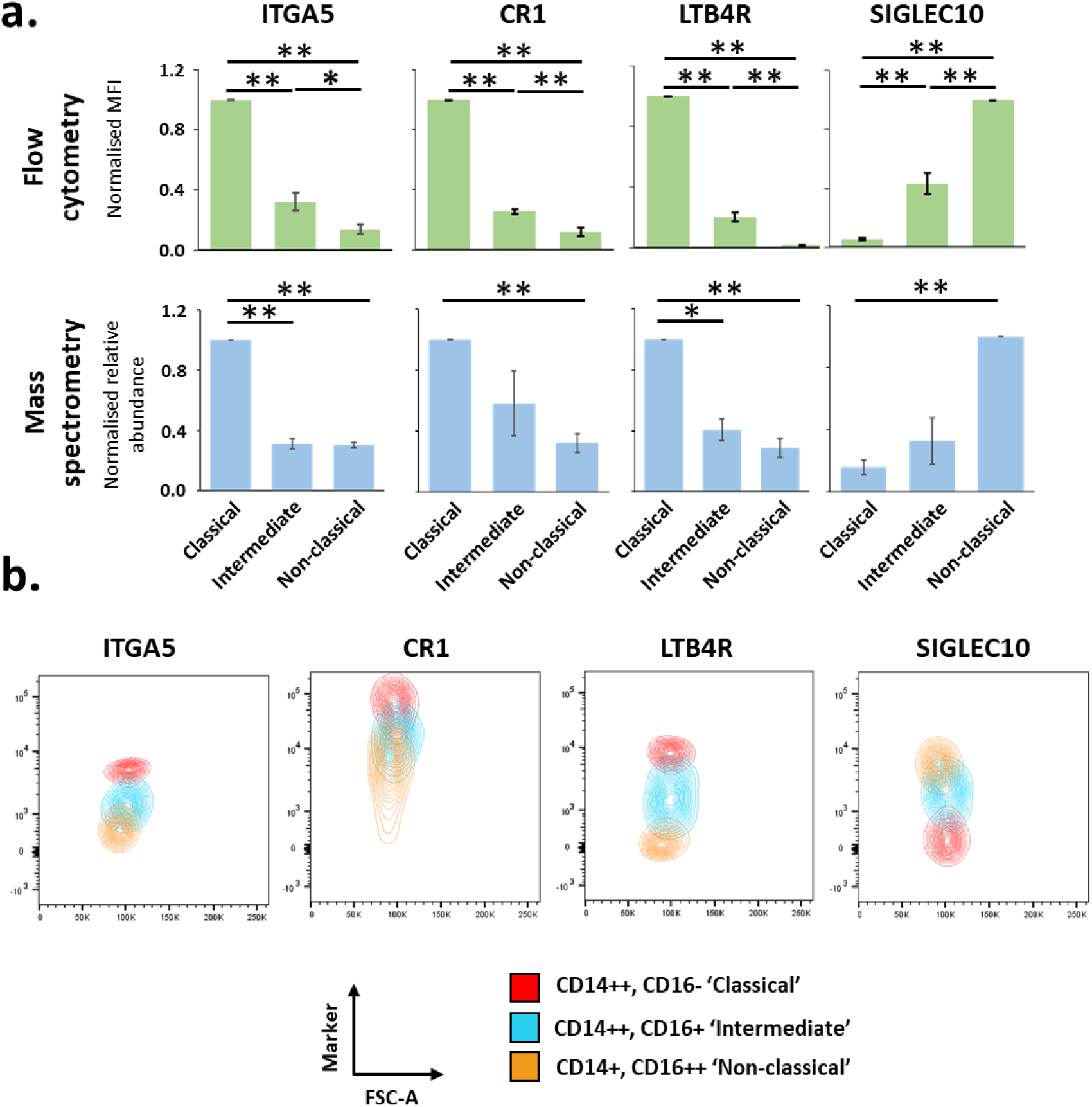
Validation of proteomics data by flow cytometry. a) Fresh human monocytes from three different donors were analysed by flow cytometry with antibodies specific to the indicated protein, in addition to anti-CD86, anti-CD14 and anti-CD16 to distinguish each monocyte subset. Corresponding profiles as determined by MS are shown in the lower panel. For both types of analyses, data were normalised to a maximum of one then averaged across the three replicates. Data are shown as mean +/- SEM. For proteomic data, a Benjamini-Hochberg-corrected two-tailed t-test was used to estimate p-values. For flow cytometry data, a two-tailed t-test was used to estimate p-values. *p< 0.05; **p<0.005. b) Representative contour plots from a single donor of flow cytometry data for each illustrated marker coloured by cell subset. FSC-A (Forward Scatter Area.) was used as an intrinsic description of cell size (which would not be expected to change between subsets) to separate the points on a second axis. This representation was advantageous compared to the alternative of histograms as it more clearly visually represents the distribution of data points with less confounding by absolute cell number. The relative proportion of each monocyte subset is different in peripheral blood, with classical monocytes predominating.

## DISCUSSION

Here we use cell surface specific protein enrichment in combination with isobaric tagging and MS3 mass spectrometry to define at high resolution the cell surface proteomes of monocyte subsets. This has not only identified novel markers of each subset but also provided a valuable resource for the future study of these crucial immune cells. This represents the first cell surface proteomic study in primary monocyte subsets. Up to now, other cell surface proteomic analyses in primary cells have been limited to CD4+ T-cells ^16^, erythrocytes ^17, 18^, NK cells, adipocytes and a subset of tumour cells^19^. We quantify 373 cell surface proteins from classical monocytes, and further quantify 313 proteins across all monocyte subsets. Based on our data we predict that the majority of the classical monocyte cell surface proteome is composed of a relatively small number of different proteins. Limitations to this study include the number of donors. To ensure all samples could be directly compared in a single TMT analysis, three subpopulations of monocytes from three donors were analysed, since only 10 different TMT tags were available. Furthermore, demographic information for each donor was not available, meaning that characteristics such as gender might bias our analysis. Nevertheless, we believe this study provides important cell surface protein level data to complement the rapidly growing array of RNA studies of different leucocyte subsets.

Here we show CR1, LTB4R and ITGA5 are markers of classical monocytes, and SIGLEC10 is a marker of non-classical monocytes. CR1 (Complement Receptor type 1 / CD35), acts as a cell surface receptor for particles opsonised by complement, facilitating their phagocytosis^20^. The specific elevation of CR1 on classical monocytes is consistent with their predominant phagocytic role.

LTB4R (also known as BLT1) is a cell surface G-protein coupled receptor for the proinflammatory leukotriene B4 (LTB4). It was originally functionally identified by subtractive cloning of cDNAs from cells differentially able to bind LTB4, and was observed to enable chemotactic response to LTB4 even in non-specialised non-immune cells^21^. Furthermore, LTB4R deletion has previously been reported to confer resistance to histopathological changes in response to a rodent model of inflammatory arthritis^22^. Given the previous link of certain monocytic subsets to inflammatory arthrititis^6^ it may be of interest to reassess the role of LTB4R positive classical monocytes in this pathology.

Integrin Alpha 5 (ITGA5, CD49e), is one of 18 mammalian alpha integrin chains which form cell surface heterodimers with integrin beta subunits^23^. Integrins are classically involved in extracellular adhesion and invasion, and also play roles in bidirectional transmembrane signalling^24^. SIGLEC10 was initially identified as a sialic acid binding protein expressed across a variety of tissues, but found at a particularly high level in organs rich in haematopoietic cells, and is notably expressed on dendritic cells and CD16+ cells ^10, 25, 26^. SIGLEC10 acts as a ligand for vascular adhesion protein 1 (VAP1), and plays a role in lymphocyte adhesion to endothelial surfaces^27^. The differential expression of both of these adhesion molecules between classical and non-classical monocytes may thus contribute to independent mobility or effector function, although future studies will be required to assess the biological function of SIGLEC10 on these cell types.

The abundance of SLC6A6 was elevated at the surface of intermediate monocytes. SLC6A6 is a known taurine and beta-alanine transporter with proposed roles in regulation of apoptosis ^28, 29^. Further studies will be required to investigate whether this molecule modulates the survival of intermediate monocytes relative to the other subsets. Similarly, our data suggested that TSPAN14 was relatively enriched on non-classical monocytes. TSPAN14 is a member of a family of tetraspanins known to regulate the subcellular localisation of metalloproteinase ADAM10^30^; this might in part explain differences in ADAM10 expression we observed between different monocyte subsets.

To determine if any particular functional class of protein was enriched in each monocyte subpopulation, we used DAVID analysis to compare proteins enriched in different subsets from our k-means analysis (**Fig. 3c**). Unfortunately, at least partly due to the relatively small background of proteins (313 proteins in total), only a single term was significantly enriched (p<0.05) within these groupings. It is therefore difficult to make confident, global statements about differences in the function of each monocyte subset. Manual inspection of the proteins enriched in each subset highlighted the presence of different cell adhesion and interaction molecules: classical – CD33, CD93, CD99; intermediate – ALCAM; non-classical – ICAM2, CD97, SELPLG (**Supplementary table S3**). We speculate that the divergent role of the monocyte subsets in varying disease states could partially be explained by their differential interactions consequent on differential expression of these molecules. For example, CD33 polymorphisms have been implicated in genetic susceptibility to Alzheimer’s disease, with risk alleles correlating with increased CD33 expression and altered phagocytic activity^31^.

As standard in our cell surface proteomic analyses we included proteins annotated with the Gene Ontology Cellular Compartment terms ‘plasma membrane’ (267/313 proteins), ‘extracellular’ (110/313 proteins), ‘cell surface’ (95/313 proteins) and a short term that indicates that the protein is membrane-integral, however that the subcellular localisation is presently unknown (7/313 proteins) (shown in Supplementary Table 1E)^13, 32^. This strategy facilitates a comprehensive coverage of the cell surface proteome. From the 313 proteins, 31 were annotated ‘extracellular’ but not ‘plasma membrane’ or ‘cell surface’. These included Lysozyme and Haptoglobin. Certain extracellular proteins can clearly bind to proteins or receptors at the plasma membrane. For example, Haptoglobin is known to bind CD163^33^, which we also detected in our analysis.

Although two previous reports have quantified the whole cell proteome of monocyte subsets^34, 35^, ours represents the first selective study of the cell surface proteome of these primary cell types. Furthermore, one of the previous studies^34^ pooled non-classical and intermediate phenotype cells prior to analysis, and the other did not examine intermediate phenotype cells^35^. We compared our data to both studies. As might be anticipated due to the low abundance and hydrophobic nature of cell surface proteins, each prior study only identified a fraction of the cell surface proteins found by selective surface biotinylation. Furthermore, for each subset of proteins previously identified as differentially expressed, there was generally poor correspondence to our data. There are a number of possible explanations for this finding, particularly including a small sample size of differentially expressed proteins that were also quantified in our study in each case. Additionally, correspondence between cell surface and whole-cell protein abundances can in any case be poor, and there may have been some confounding in one of the prior studies from isolation and measurement of protein samples in five different locations, using four different types of mass spectrometer^35^. Neither prior study examined intermediate-phenotype cells. We believe this shows the clear benefits of selective interrogation of the cell surface proteome, in a single multiplexed experiment that can make precise quantitative comparisons.

Although the monocyte subsets can be delineated using CD14/CD16 expression, use of further markers in parallel could aid distinction of subgroups which may appear as a near continuum if viewed in only two dimensions^12^. It has previously been suggested that the intermediate monocyte subset may be a transitional state between the two larger circulating monocyte populations, and the classical subset are more distinct from intermediate and non-classical cells than the latter two populations are from one another. Our data support this second observation. However, we also identified proteins that are uniquely up- or down-regulated in intermediate cells, arguing against the hypothesis that these cells may be purely transitional. Future work will be required to determine which combinations of markers can be used to definitively separate individual subsets, and will expand recent single cell transcriptomic data which has suggested that intermediate monocytes are more heterogenous than previously anticipated^9, 36^. Additionally, further studies will also be required to examine the monocyte cell surface from individuals with diseases including asthma, rheumatoid arthritis and sarcoidosis to determine whether subpopulations exhibit phenotypic differences in these conditions. We thus provide an orthogonal analysis of monocyte subgroups to complement previous transcriptomic studies which will facilitate many future studies of monocytes. This valuable resource may also be useful to assist elucidation of the true nature of intermediate monocytes.

## MATERIALS AND METHODS

### Monocyte isolation

For proteomic studies, leukocyte enriched blood samples were obtained from healthy UK-based blood donors via NHS blood and transplant (NHSBT, Cambridge, UK). No further information about these donors was available from NHSBT, in line with their conditions of supply. Cells were eluted and PBMC isolated by centrifugation on a Ficoll gradient (GE Healthcare) for 20 minutes at 800g. PBMC were aspirated from the interface, and monocytes isolated using a negative selection kit (‘Pan Monocyte Enrichment’, Miltenyi Biotec, 130-096-537). Monocyte subsets were subsequently defined by flow cytometry after staining with anti-CD14, anti-CD16 and anti-CD86. For fluorescence activated cell sorting on either Influx or Aria III cell sorter (Becton Dickinson), gating strategies are shown in Fig. 1. Live cells were defined by a forward and side-scatter gate. Live CD86+ cells were sorted into three subsets defined by CD14 and CD16 expression. Samples used for flow-cytometry based phenotypic validation of cell surface marker expression (Fig. 4) were derived from healthy donors of European ancestry local to Cambridge, UK. Here, 5-10ml of whole anticoagulated blood was collected from each donor and subjected to enrichment and staining as described above, including stains for anti-CD14, anti-CD16 and anti-CD86 in addition to individual markers of interest. Donated blood was collected with informed consent in accordance with the Declaration of Helsinki. Ethical approval was obtained from University of Cambridge Human Biology Research Ethics Committee (HBREC.2016.011).

### Plasma membrane profiling

Plasma membrane profiling was performed broadly as previously described^13^, with the following modifications. The total number of monocytes biotinylated was dependent on the yield from enrichment procedures detailed above, typically in the range of 10^5^-10^6^ cells for intermediate and non-classical monocytes, and 10^7^ for classical monocytes. Precise sorted counts were collected during each enrichment from the Influx or Aria III cell sorter. Each of the three donated samples was biotinylated and labelled separately, combining at the final stage prior to TMT analysis. Briefly, for each enriched monocyte subset sample, sialic acid residues at the cell surface were oxidized using sodium meta-periodate (Thermo) and biotinylated with aminooxy-biotin (Biotium). Following quenching, cells were incubated in 1% (v/v) Triton X-100 lysis buffer (10mM Tris HCl, 1.6% Triton, 150mM NaCl). Biotinylated glycoproteins were precipitated using high affinity streptavidin agarose beads (Pierce), and washed extensively. Captured protein was then reduced with dithiothreitol (DTT), alkylated with iodoacetamide (Sigma) and digested on-bead with trypsin (Promega) in 200 mM HEPES pH 8.5 for 3 hours. Trypsin cleaves C-terminal to basic residues, except when they are N-terminal to a Proline residue. Tryptic peptides were collected and the whole sample labelled using TMT reagents following dilution in 200mM HEPES pH 8.5 adjusted to a final 30% acetonitrile concentration (v/v). To ensure analysis of each cell type in a 1:1:1 ratio, a fraction of each labelled peptide sample was combined in proportion to the number of cells collected for each subset. Labelling was as follows; classical monocytes donation 1 (126), intermediate monocytes donation 1 (127N), non-classical monocytes donation 1(127C), classical monocytes donation 2 (128N), intermediate monocytes donation 2 (128C), non-classical monocytes donation 2 (129N), classical monocytes donation 3 (129C), intermediate monocytes donation 3 (130N), non-classical monocytes donation 3 (130C). The reaction was quenched with hydroxylamine, and TMT-labelled samples combined in a 1:1:1:1:1:1:1:1:1 (all subsets) or 1:1:1 (Classical only) ratio. Labelled peptides were subjected to C18 solid-phase extraction (Sep-Pak, Waters) and vacuum-centrifuged to near-dryness.

### Offline SCX fractionation

Offline fractionation was performed as previously described^14, 37^ with the following modifications. 10 mg of PolySulfethyl A bulk material (Nest Group Inc) was loaded on to a fritted 200 μl tip in 100% Methanol using a vacuum manifold. SCX material was conditioned slowly with 1ml SCX buffer A (7 mM KH_2_PO_4_, pH 2.65, 30% Acetonitrile), then 0.5 ml SCX buffer B (7 mM KH_2_PO_4_, pH 2.65, 350mM KCl, 30% Acetonitrile) then 2ml SCX buffer A. Dried peptides were resuspended in 500 μl SCX buffer A and added to the tip at a flow rate of ∼150 μl/min, followed by a 150 μl wash with SCX buffer A. Fractions were eluted in 150 μl buffer at increasing K+ concentrations (10, 25, 40, 60, 90, 150 mM KCl), vacuum-centrifuged to near dryness then desalted using StageTips.

### LC-MS3

LC-MS3 was performed with modifications as previously described^38, 39^. Mass spectrometry data was acquired using an Orbitrap Lumos (Thermo Fisher Scientific, San Jose, CA). An Ultimate 3000 RSLC nano UHPLC equipped with a 300 µm ID x 5 mm Acclaim PepMap µ-Precolumn (Thermo Fisher Scientific) and a 75 µm ID x 50 cm 2.1 µm particle Acclaim PepMap RSLC analytical column was used. An unfractionated singleshot was analysed initially to ensure similar peptide loading across each TMT channel, thus avoiding the need for excessive electronic normalization. As all normalisation factors were >0.33 and <3.0, data for each singleshot experiment was analysed with data for the corresponding fractions to increase the overall number of peptides quantified. For monocyte subset analysis, 2µl of fractionated peptide was initially analysed by mass spectrometry, followed by the remainder of the sample having observed satisfactory chromatography.

For LC/MS3, loading solvent was 0.1% trifluoroacetic acid (TFA), analytical solvent A: 0.1% FA and B: 80% acetonitrile + 0.1% FA. All separations were carried out at 55°C. Samples were loaded at 10 µl/minute for 5 minutes in loading solvent before beginning the analytical gradient. The following gradient was used: 3-7% B over 4 minutes, 7-37% B over 58 minutes, 37-95% B over 4 minutes followed by a 2 minute wash at 95% B and equilibration at 3% B for 15 minutes. Each analysis used a MultiNotch MS3-based TMT method ^40, 41^. The following settings were used: MS1: 380-1500 Th, Quadrupole isolation, 120,000 Resolution and 2×10^5^ AGC target, 50 ms maximum injection time. MS2: Quadrupole isolation at an isolation width of m/z 0.7, CID fragmentation (NCE 35) with ion trap scanning out in rapid mode from m/z 120, 1.5×10^4^ AGC target, 300 ms maximum injection time in centroid mode. MS3: in Synchronous Precursor Selection mode the top 10 MS2 ions were selected for HCD fragmentation (NCE 65) and scanned in the Orbitrap at 60,000 resolution with an AGC target of 1.5×10^5^ and a maximum accumulation time of 250 ms, ions were not accumulated for all parallelisable time. The entire MS/MS/MS cycle had a target time of 3 s. Dynamic exclusion was set to +/-10 ppm for 70 s. MS2 fragmentation was trigged on precursors 5×10^3^ counts and above.

### Data analysis

Data analysis was performed with modifications as previously described^18, 38^. In the following description, we list the first report in the literature for each relevant algorithm. Mass spectra were processed using a Sequest-based software pipeline for quantitative proteomics, “MassPike”, through a collaborative arrangement with Professor Steve Gygi’s laboratory at Harvard Medical School. MS spectra were converted to mzXML using an extractor built upon Thermo Fisher’s RAW File Reader library (version 4.0.26). In this extractor, the standard mzXML format has been augmented with additional custom fields that are specific to ion trap and Orbitrap mass spectrometry and essential for TMT quantitation. These additional fields include ion injection times for each scan, Fourier Transform-derived baseline and noise values calculated for every Orbitrap scan, isolation widths for each scan type, scan event numbers, and elapsed scan times. This software is a component of the MassPike software platform and is licensed by Harvard Medical School.

A combined database was constructed from the human Uniprot database (26^th^ January, 2017), and common contaminants such as porcine trypsin. The combined database was concatenated with a reverse database composed of all protein sequences in reversed order. Searches were performed using a 20 ppm precursor ion tolerance^42^. Fragment ion tolerance was set to 1 Da. TMT tags on lysine residues and peptide N termini (229.162932 Da) and carbamidomethylation of cysteine residues (57.02146 Da) were set as static modifications, while oxidation of methionine residues (15.99492 Da) was set as a variable modification.

To control the fraction of erroneous protein identifications, a target-decoy strategy was employed ^43, 44^. Peptide spectral matches (PSMs) were filtered to an initial peptide-level false discovery rate (FDR) of 1% with subsequent filtering to attain a final protein-level FDR of 1% ^45, 46^. PSM filtering was performed using a linear discriminant analysis, as described previously ^47^. This distinguishes correct from incorrect peptide IDs in a manner analogous to the widely used Percolator algorithm ^48^, though employing a distinct machine learning algorithm. The following parameters were considered: XCorr (minimum 1), ΔCn, missed cleavages, peptide length, charge state, and precursor mass accuracy. Protein assembly was guided by principles of parsimony to produce the smallest set of proteins necessary to account for all observed peptides ^47^.

Proteins were quantified by summing TMT reporter ion counts across all matching peptide-spectral matches using “MassPike”, as described previously ^40, 41^. Briefly, a 0.003 Th window around the theoretical m/z of each reporter ion (126, 127n, 127c, 128n, 128c, 129n, 129c, 130n, 130c) was scanned for ions, and the maximum intensity nearest to the theoretical m/z was used. The primary determinant of quantitation quality is the number of TMT reporter ions detected in each MS3 spectrum, which is directly proportional to the signal-to-noise (S:N) ratio observed for each ion ^49^. Conservatively, every individual peptide used for quantitation was required to contribute sufficient TMT reporter ions so that each on its own could be expected to provide a representative picture of relative protein abundance ^40^. An isolation specificity filter was additionally employed to minimise peptide co-isolation ^50^. Peptide-spectral matches with poor quality MS3 spectra (more than 9 TMT channels missing and/or a combined S:N ratio of less than 135 (9-plex, monocyte subsets) or 45 (3-plex, classical monocytes) across all TMT reporter ions) or no MS3 spectra at all were excluded from quantitation. Peptides meeting the stated criteria for reliable quantitation were then summed by parent protein, in effect weighting the contributions of individual peptides to the total protein signal based on their individual TMT reporter ion yields. Protein quantitation values were exported for further analysis in Microsoft Excel.

For protein quantitation, reverse and contaminant proteins were removed. For further analysis and display in figures, fractional TMT signals were used (i.e. reporting the fraction of maximal signal observed for each protein in each TMT channel, rather than the absolute normalized signal intensity). This effectively corrected for differences in the numbers of peptides observed per protein.

Proteins were filtered to include those most likely to be present at the cell surface with high confidence. These contained proteins with the Gene Ontology (GO) terms of ‘plasma membrane’ (PM), ‘cell surface’ (CS), ‘extracellular’ (XC) or with a short 4- or 5-part GO cellular compartment term that included ‘integral to membrane’, but with no subcellular assignment ^13^ (ShG).

To estimate the relative abundance of each protein, a method based on iBAQ was employed. The summed MS1 maximum precursor intensity for each protein across all matching peptides was calculated. Each value was divided by the number of theoretically observable tryptic peptides 7-30 amino acids in length for the respective protein, as determined by *in silico* trypsin digestion of human Swissprot canonical and isoform database (2017_01_26) using the OrgMassSpecR^51^ package in R 3.5.1^52^.

For Fig. 1b, the Database for Annotation, Visualisation and Integrated Discovery (DAVID) version 6.8, was used to determine pathway enrichment^53^. 373 proteins identified at the plasma membrane of classical monocytes were searched against a background of the whole human proteome.

Hierarchical centroid clustering based on Euclidian distance or uncentered correlation was performed using Cluster 3.0^54^ (Stanford University) and visualised using Java Treeview^55^ (http://jtreeview.sourceforge.net). Principle component analysis and K-means analysis to determine the number of distinct patterns of protein expression between monocyte subsets was performed using XLSTAT v2019.1.2 (Addinsoft). Once the number of expression profiles was determined, Cluster 3.0 was used to perform K-means clustering.

### Flow cytometric analysis of monocyte surface proteins

Monocytes were obtained by negative selection using a Pan-Monocyte enrichment kit as described above. Cells were incubated with Fc blocking reagent (Biolegend Human Trustain FcX) for 15 min at 4°C. Four-colour staining was then used to validate phenotype, using anti-CD14, anti-CD16, anti-CD86 and an antibody against the marker of interest for 15 min at 4°C. Specific antibodies used were: APC-Cy7 anti-CD14 (Biolegend, 325620), PE anti-CD14 (Biolegend, 301850), BV421 anti-CD16 (Becton Dickinson, 562874), PE-Cy7 anti-CD86 (Becton Dickinson, 561128), FITC anti-ITGA5 (Miltenyi Biotec, 130-110-592), APC anti-SIGLEC10 (Miltenyi Biotec, 130-103-731), PE anti-LTB4R (BioRad, MCA2108PET), PE anti-CD35 (Miltenyi Biotec, 130-099-913). Cells were washed in PBS/0.4% (v/v) citrate/0.5% (v/v) BSA prior to fixation and analysis on a BD LSR Fortessa (Beckton Dickinson). After gating by forward and side-scatter, ≥30,000 events were captured. Data was analysed using FlowJo V10 (FlowJo, LLC). Monocytes were defined using forward and side-scatter filters and positive CD86 expression as shown in Supplementary Fig. S1.

For Fig. 4, monocytes were gated into subsets: CD14++, CD16- (Classical), CD14++, CD16+ (Intermediate), CD14+, CD16++ (Non-classical). MFI values for each subset were normalised to a maximum of 1 prior to averaging and calculation or the standard deviation and standard error of the mean.

### Experimental design

For each proteomic analysis (classical monocyte or subset analysis) three independent samples were obtained and prepared separately prior to simultaneous isobaric tag labelling and further analysis. Triplicate analysis was chosen to take advantage of the capability to study up to 10 samples simultaneously using tandem mass tags. As such, three complete sets of three subsets were chosen. Validation studies using flow cytometry were performed in biological triplicate from three independent donors as described above. Flow cytometry fluorophore staining was performed with singly stained and unstained control populations in addition to the test samples. For proteomic data, a Benjamini-Hochberg-corrected two-tailed t-test was used to estimate p-values. For flow cytometry data, a two-tailed t-test was used to estimate p-values based on an observed near-normal distribution of signal within subgroups.

### Data availability

The mass spectrometry proteomics data has been deposited to the ProteomeXchange Consortium (http://www.proteomexchange.org/) via the PRIDE^56^ partner repository, project identifier PXD013832. Project name ‘Comprehensive cell surface proteomics defines markers of classical, intermediate and non-classical monocytes’.

## ACKNOWLEDGEMENTS

We are grateful to Prof. Steven Gygi for providing access to the “MassPike” software pipeline for quantitative proteomics. This work was supported by an Evelyn Trust research training fellowship (18/27) to BJR, a Wellcome Trust Senior Clinical Research Fellowship (108070/Z/15/Z) to MPW and a strategic award to Cambridge Institute for Medical Research from the Wellcome Trust (100140). This study was additionally supported by the Cambridge Biomedical Research Centre, UK. We would like to thank the NIHR Cambridge BRC phenotyping hub for their assistance in sorting primary monocytes into their subgroups.

## AUTHOR CONTRIBUTIONS

BJR and MPW designed and performed the experiments. BJR and LS isolated monocytes from leukocyte enriched donor samples. RA collected proteomic data. BJR, JH and MPW analysed proteomic data. BJR and MPW wrote the manuscript. MPW supervised all research.

