## Supplementary information for "Comprehensive cell surface proteomics defines markers of classical, intermediate and non-classical monocytes"

**Affiliations:**

### Supplementary Information

**Supplementary table 1**

- **S1A** – Unmodified and unfiltered signal:noise values for classical monocyte samples, filtered by a signal:noise threshold of 45.
- **S1B** - Unmodified and unfiltered signal:noise values for monocyte subsets samples, filtered by a signal:noise threshold of 135
- **S1C** – Classical monocyte and monocyte subset signal:noise data, following normalisation and summation of samples.
- **S1D** – Classical monocyte data filtered by GO term (NB. HLA molecules not combined)
- **S1E** – Monocyte subset data filtered by GO term (NB. HLA molecules not combined)
- **S1F** – An interactive worksheet of the data contained within the spreadsheet. Enter the gene name of interest into the yellow box to view summarised data.

**Supplementary table 2**

- **S2A** - Unmodified and unfiltered signal:noise values for peptides for data filtered by a signal:noise threshold of 45.
- **S2B** - Unmodified and unfiltered signal:noise values for peptides for data filtered by a signal:noise threshold of 135.
- **S2C** – Protein groups, and peptide assignment to these groups

**Supplementary table 3**

- A summary of the reported functions of proteins mentioned by name in the manuscript. The references are included as PubMed IDs (PMID).

### Supplementary Figure S1

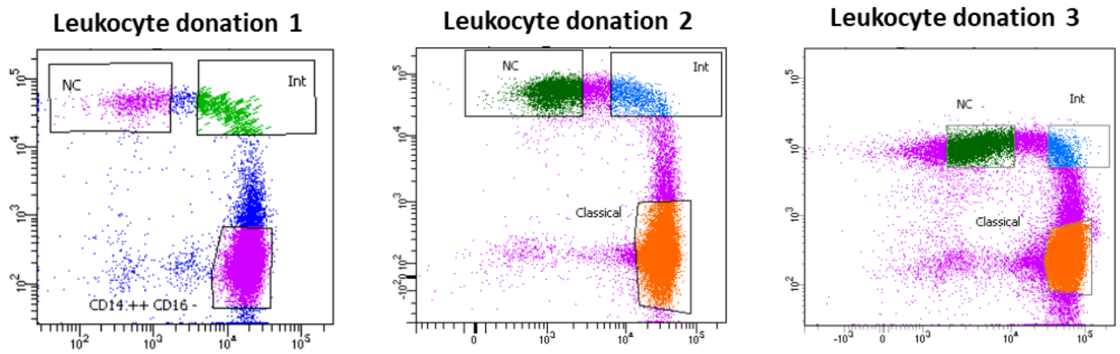


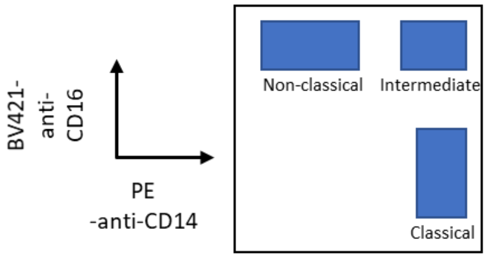


| | Sort 1 | | Sort 2 | | Sort 3 | |
|---|---|---|---|---|---|---|
| | % Pre-sort purity | % Post-sort purity | % Pre-sort purity | % Post-sort purity | % Pre-sort purity | % Post-sort purity |
| Classical | 87.2 | 98.6 | 92.4 | 99.9 | 78.7 | 99.5 |
| Intermediate | 2.1 | 99.1 | 3.72 | 92.4 | 2.61 | 99.3 |
| Non-classical | 4.5 | 100 | 14.9 | 99.5 | 6.77 | 96.4 |

**Supplementary Figure S1 – Fluorescence activated cell sorting of each donation used in this study.** Original CD14 vs CD16 dot-plots of the three sorts used in this study with gating boundaries. Below is a schematic of the cell subset enriched by each gate and tables of pre- and post- sort purities following FACS for each leukocyte donation.

### Supplementary Figure S2

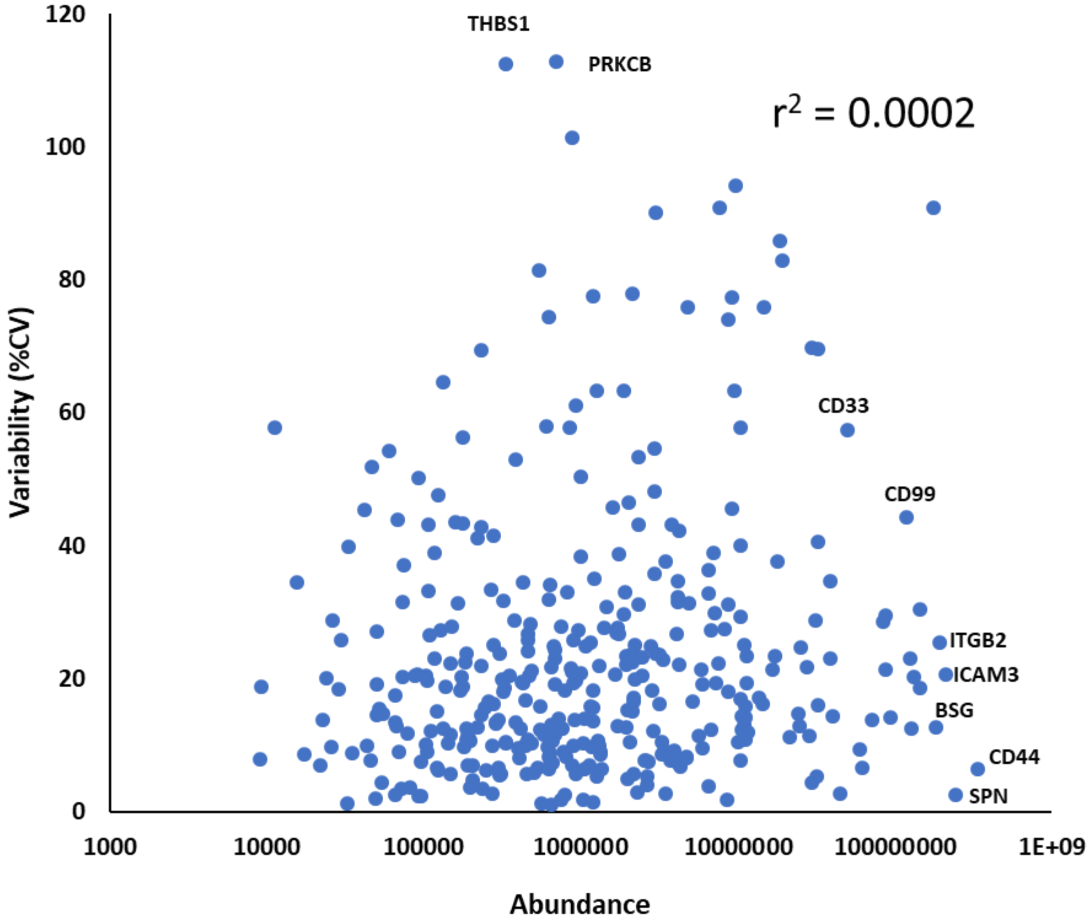


**Supplementary Figure S2 – Abundance of classical monocyte cell surface proteins did not correlate with calculated variability across three samples**. Coefficient of variation (%CV) for each protein across each of three samples was plotted against an estimate of protein abundance derived using an adapted iBAQ method as described in the text. HLA molecules are represented individually. The $r^2$ coefficient of determination was 0.0002.

#### Supplementary Figure S3

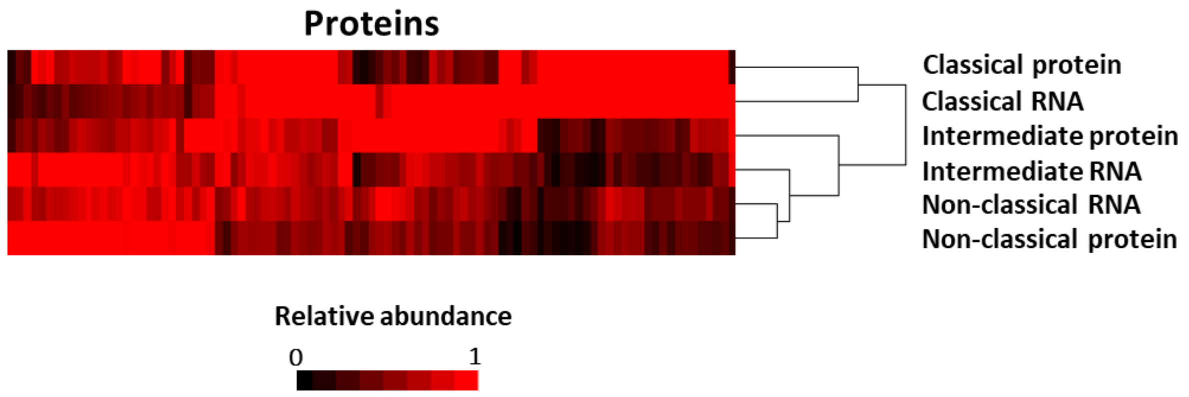


**Supplementary Figure S3 – Comparison of proteomics to published RNA microarray data.** Hierarchical cluster analysis of 95 proteins and transcripts. For each of three replicates, signal:noise for each protein was averaged then normalised to a maximum of one. A similar strategy was employed for a comprehensive transcriptomic dataset (Wong et al.).

### Supplementary Figure S4

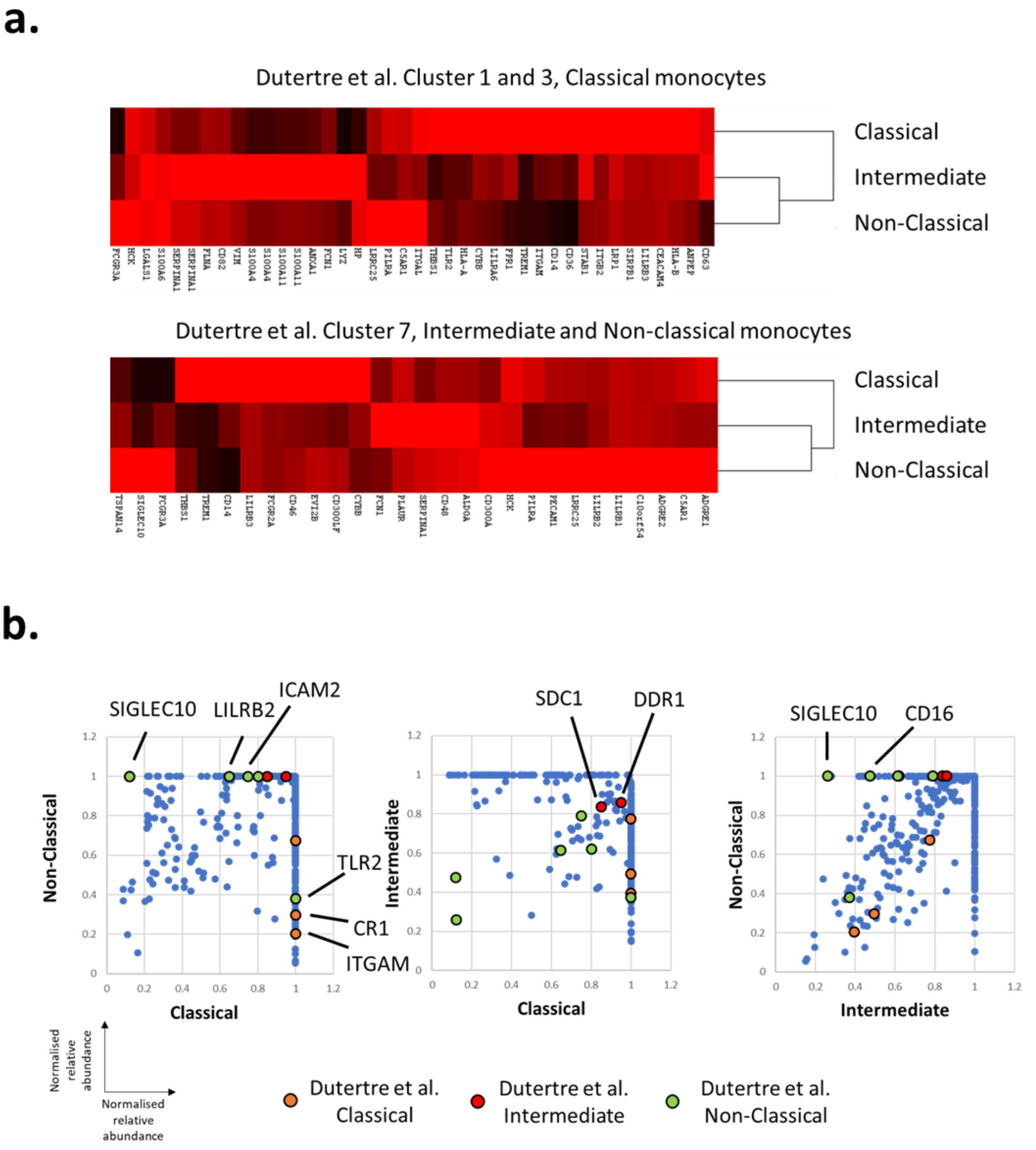


**Supplementary Figure S4 – Comparison of proteomics to published single cell RNA-seq data.**

(a) Hierarchical cluster analysis of cell surface protein abundance for differentially expressed genes identified by Dutertre at al. DEGs for CD14$^{hi}$CD16$^{-}$ (classical) and CD16$^{+}$ (intermediate and non-classical) monocytes corresponded well to discriminating proteins in our data. Particularly for the upper panel, the majority of 'classical' DEGs were either strongly positively or negatively expressed at the cell surface of classical monocytes. (b) Comparison of the monocyte surface proteome to differentially expressed proteins (DEPs) determined from flow cytometry by Dutertre et al. For each protein, TMT signal:noise data (Supplementary table S1e) were averaged for each monocyte subset, before normalising to a maximum of one. Three pairwise comparisons are shown with DEPs highlighted. For classical compared to non-classical monocytes, the markers were highly discriminatory (all DEPs lay at extremes away from the y=x line, as would be expected for differentially expressed proteins). The least convincing were the intermediate monocyte markers, of which only two (SDC1, DDR1) were present in both data sets. These exhibited similar expression (near the (1,1) point) on classical and non-classical cells. This observation is consistent with the flow cytometry data from Dutertre et al. where for these markers the difference in staining for intermediate monocytes compared to the other monocyte subsets was small. This exemplifies the difficulty of identifying intermediate monocyte-specific cell surface markers. The only major disparity between the datasets was TLR2/CD282, which Dutertre et al. suggested to be most highly expressed in non-classical monocytes. In comparison, we found this marker to be most abundant on classical monocytes. Further work will be required to assess whether TLR2 expression varies between larger cohorts of individual donors.
